# Tracking progress towards malaria elimination in China: estimates of reproduction numbers and their spatiotemporal variation

**DOI:** 10.1101/628842

**Authors:** Isobel Routledge, Shengjie Lai, Katherine E Battle, Azra C Ghani, Manuel Gomez-Rodriguez, Kyle B Gustafson, Swapnil Mishra, Joshua L Proctor, Andrew J Tatem, Zhongjie Li, Samir Bhatt

## Abstract

China reported zero locally-acquired malaria cases in 2017 and 2018. Understanding the spatio-temporal pattern underlying this decline, especially the relationship between locally-acquired and imported cases, can inform efforts to maintain elimination and prevent re-emergence. This is particularly pertinent in Yunnan province, where the potential for local transmission is highest. Using a geo-located individual-level dataset of cases recorded in Yunnan province between 2011 and 2016, we jointly estimate the case reproduction number, *R*_*c*_, and the number of unobserved sources of infection. We use these estimates within spatio-temporal geostatistical models to map how transmission varied over time and space, estimate the timeline to elimination and the risk of resurgence. Our estimates suggest that, maintaining current intervention efforts, Yunnan is unlikely to experience sustained local transmission up to 2020. However, even with a mean *R*_*c*_ of 0.005 projected for the year 2019, locally-acquired cases are possible due to high levels of importation.

---

In 2016 the World Health Organisation listed 21 countries for whom it would be realistic to achieve elimination of malaria by 2020, defined as zero indigenous cases over three consecutive years^1^. The largest of these is the People’s Republic of China (thereafter called China). In 2017 China reported no indigenous malaria cases for the first time since malaria became a notifiable disease in 1956^2,3^. The country has experienced a major decline in the burden of malaria, from an annual incidence of 24 million cases (2961 cases per 100,000) in 1970^4^. This reduction has been attributed to a combination of socioeconomic improvements and the scale-up of interventions to control malaria^5^. In 2010, China set out an ambitious plan for the national elimination of malaria by 2020 (the National Malaria Elimination Programme, NMEP). Elements of the plan included improved surveillance, timely response, more effective and sensitive risk assessment tools and improved diagnostics^6^. A key policy change implemented in 2010 as part of the NMEP was the introduction of the 1-3-7 system: aiming for case reporting in one day, which is then investigated within three days, with a focused investigation and action taken in under seven days^7^.

Although China is making rapid progress towards this goal, 2,675 imported cases were reported in 2017, highlighting the risk of re-introduction^3^. Large numbers of people move between China and malaria endemic countries, both from sub-Saharan Africa and from South East Asia ^8,9^, driven by tourism and Chinese oversea investment^10^. Concerns remain about re-emergence of malaria, which has occurred several times in the early 2000s as a result of importation and favourable climatic conditions for competent vectors^11^. Therefore, in order to achieve three consecutive years of zero indigenous cases (the requirement for WHO certification of elimination), a sustained and targeted investment in surveillance together with efficient treatment is necessary.

Yunnan province has recorded malaria outbreaks and remains an identified foci of residual transmission as other areas in the country have reached elimination^12–16^. The province shares borders with Myanmar, Vietnam and Laos and has a strong agricultural focus. Previous studies suggest that seasonal agricultural workers and farmers are at highest risk of contracting malaria in Yunnan, with rice yield and the proportion of rural employees being spatial factors positively associated with malaria incidence^17^. The border region of Myanmar and Yunnan is generally ecologically suitable for malaria transmission, has a large mobile population, with few natural geographic borders separating the two countries, as well as being a site of socio-political conflict and instability^18^. In this context, it can be unclear whether there is any sustained local transmission or if all the observed cases are the result of short, stuttering transmission chains following importation into suitable areas. As the area of highest concern for re-emergence in China and the last to reach zero cases, we therefore sought to characterise the transmission dynamics of both *Plasmodium vivax* and *Plasmodium falciparum* in the region as China approaches elimination certification.

Methods from outbreak analysis and network research have recently been developed and applied to quantify the transmission of malaria and other infectious diseases in near-elimination and epidemic settings^19–21^. In near elimination contexts with strong surveillance systems, traditional metrics of malaria such as parasite prevalence are not appropriate due to small numbers and extremely sparse and spatiotemporally heterogeneous distributions of infections. However due to the strength of the surveillance system in China, detailed information is available about each individual case (including the time of symptom onset and location of residence), and case reporting is believed to be very high. By adapting and applying a diffusion network approach ^22^ within a Bayesian framework, we quantify case reproduction numbers, *R*_*c*_, and uncertainty in these estimates for all *P. vivax* and *P. falciparum* cases of malaria recorded in Yunnan province between 2011 and 2016. We incorporate these estimates into geostatistical models and time series analysis to estimate how *R*_*c*_ varied over space and time which we use to estimate timelines to elimination and likelihood of resurgence.

## Results

### *R*_*c*_ estimates over time

Between 2011 and 2016, 3496 cases of probable and confirmed *P. vivax* infection including mixed infections were observed in Yunnan province (2881 imported, 615 locally acquired). Including mixed infections, 818 *P. falciparum* infections were observed, of which 75 were locally acquired. The mean *R*_*c*_ value estimated for *P. vivax* during this period was 0.171 (95% CI = 0.165, 0.178) and 0.089 (95% CI = 0.076, 0.103) for *P. falciparum* cases (Supplementary Figure 1). We estimate a decline in *R*_*c*_ over time for both *P. vivax* (Figure 1A and 1B) and *P. falciparum* (Figure 1C and 1D), with the most rapid declines occurring between 2012 and 2014 (Figure 1A and 1C). No *R*_*c*_ values above one were observed after 2014 for either species. These findings are consistent with varying levels of uncertainty about the serial interval distribution (Supplementary Figures 2 and 3).

**Figure 1:**
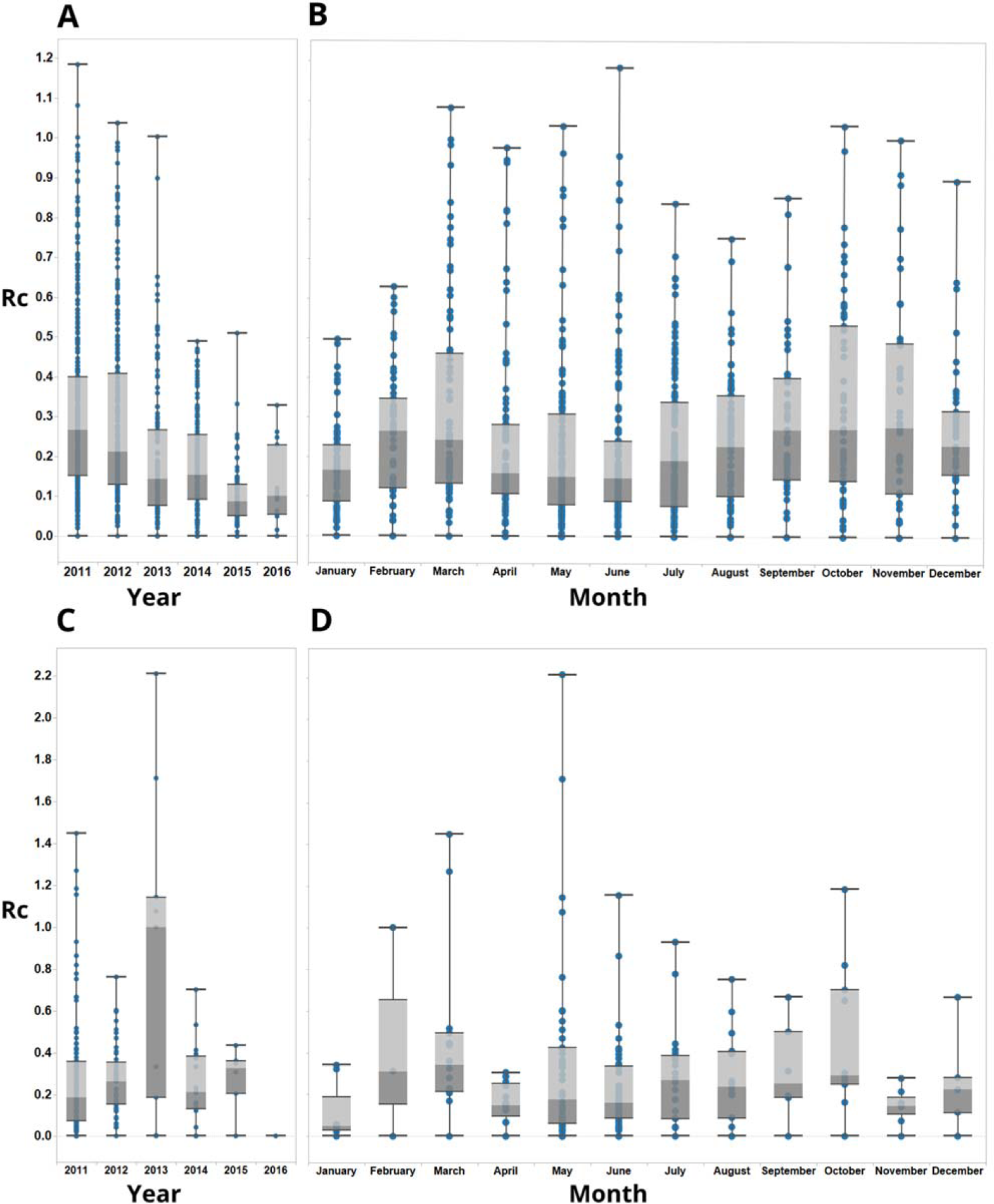
Boxplots showing *R*_*c*_ estimates for *P. vivax* (A and B) and *P. falciparum* (C and D), aggregated by year (A and C) and month (B and D) of symptom onset. Points represent individual *R*_*c*_ estimates. Boxplots show median, upper and lower quartiles for *R*_*c*_ each.

**Figure 2:**
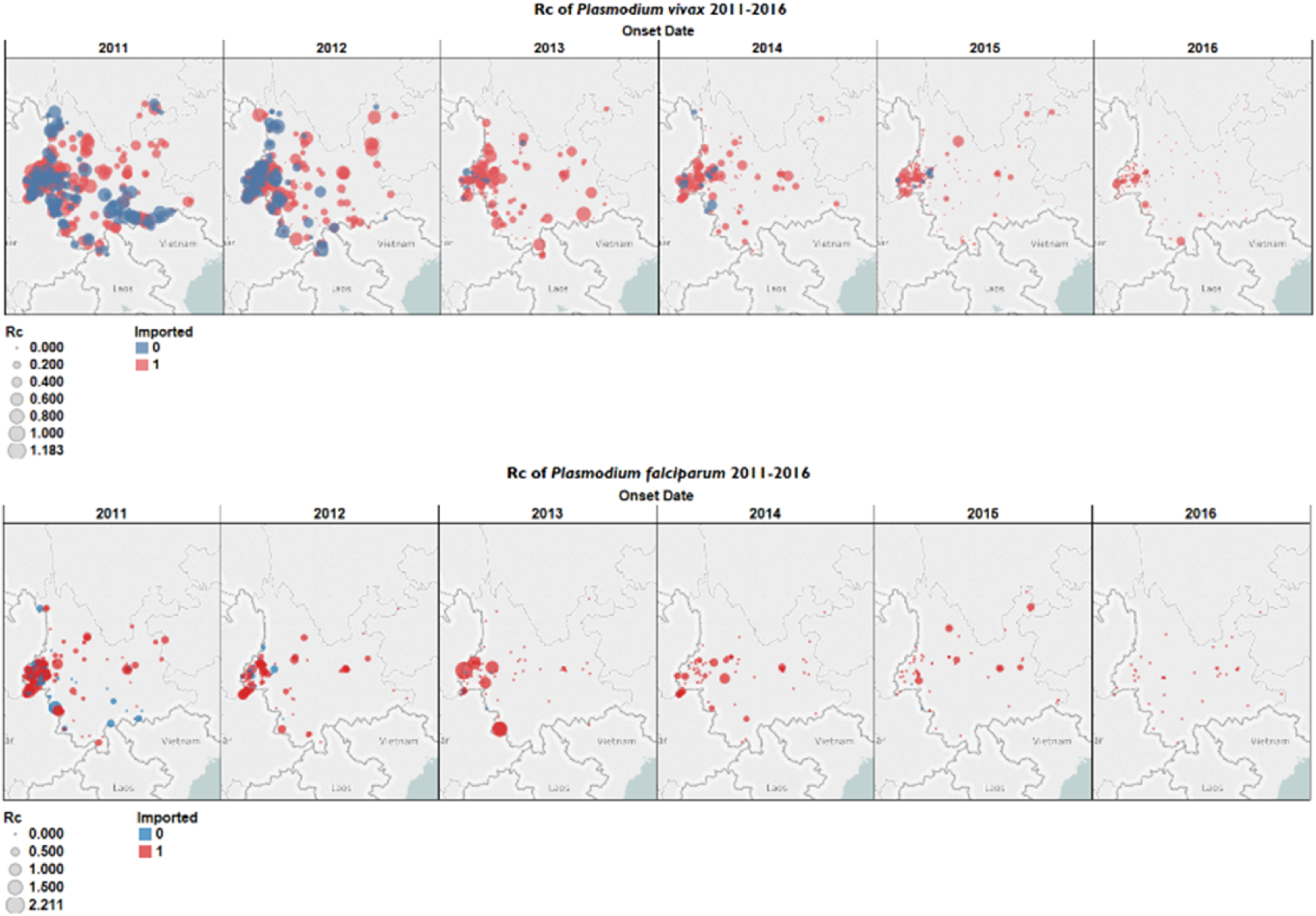
Map of *R*_*c*_ estimates by year for A) *P. vivax* and B) *P. falciparum*. Blue points represent locally acquired cases; red points represent imported cases. The diameter of the point represents the size of the *R*_*c*_ estimate.

**Figure 3:**
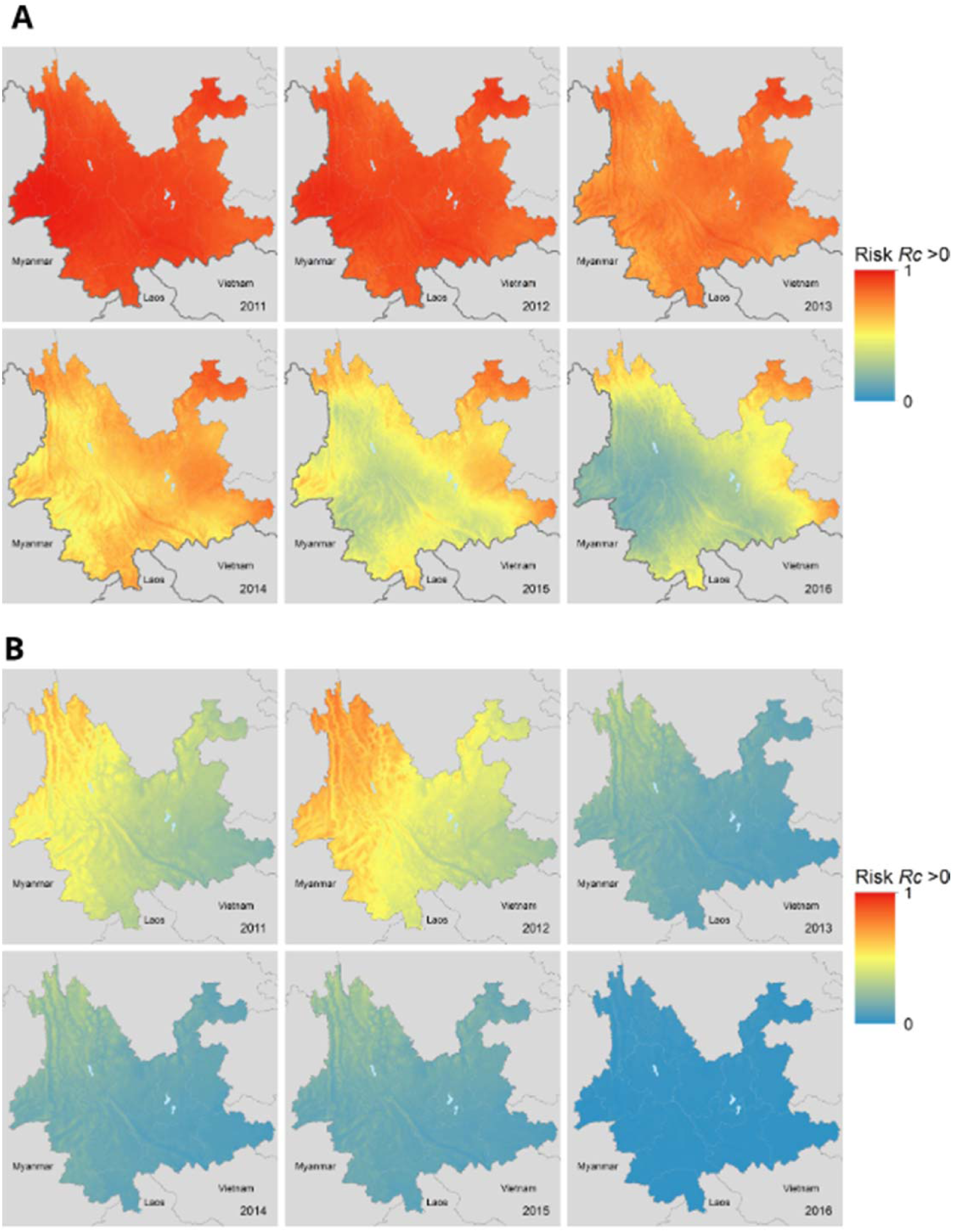
Map of risk of *R*_*c*_ > 0 and uncertainty in this estimate for A) *P. vivax* and B) *P. falciparum* malaria across Yunnan province in each year 2011-2016.

### Unobserved sources of infection

For *P. vivax*, 19 out of 615 locally acquired cases were estimated to have a moderate chance of having an unobserved source of infection (estimated 0.8 ≥ *ε* ≥ 0.5) and 2 cases were estimated to have a high chance of an unobserved source of infection (estimated *ε* ≥ 0.8). Together, this represents 3% of locally acquired cases with a moderate to high chance of external infection sources. For *P. falciparum*, 2 out of 75 local cases were estimated to have a high chance of having an unobserved source of infection (estimated *ε* ≥ 0.8) and no other cases were estimated to have a moderate change of having an unobserved source of infection (Supplementary Figure 5).

### Spatial patterns of *R*_*c*_

As transmission declined between 2011 and 2016, we observed a reduction in the incidence of locally-acquired cases which is reflected in a reduction in our estimates of the reproduction number of each locally-acquired case for both species and with a more focal spatial distribution of cases (Figure 2A and 2B). We estimate a decline in the probability of a reproduction number for a *P. vivax* case being above zero over this period (Figure 3A and 3B), with the central parts of the province being the first to reach lower risks of non-zero *R*_*c*_. The border area neighbouring Myanmar, where most cases were observed, had the lowest amount of uncertainty in the estimates. *P. falciparum* shows a decline in risk of *R*_*c*_ > 0 across the province, with the more isolated areas in the north of the province showing both the highest predicted risk but also the most uncertainty, due to a lack of cases observed there (Supplementary Figure 6). By 2016 all areas have reached a low risk, although there is more uncertainty in these estimates compared to *P. vivax*, almost certainly due to the smaller sample size.

### Short – term predictions and temporal patterns in timeseries of *Plasmodium vivax* cases

Using a time series method to make short-term predictions, we estimate a posterior mean *R*_*c*_ of 0.005 (95% CI = 0 – 0.34) for *Plasmodium vivax* cases in 2019 (Figure 4A). We observe and overall declining trend, with the fitted trend for *R*_*c*_ (which estimates the general trend, separate to the influence of seasonal and holiday effects) declining from 0.31 (95% CI = 0.31, 0.34) at the start of 2011 to 0.004 (95% CI =0.002-0.006) by the end of 2019 (Figure 4B). We estimate a small effect of holiday periods to differences in *R*_*c*_ observed, with Chinese New Year and National Day associated with small increase risk in *R*_*c*_ of 16% (95% CI = −112%, 152%) and 39% (95% CI = −43%, 118%) (Figure 4B) which in this very low transmission context could increase the probability of small outbreaks of local transmission in areas in which high rates of importation occur, although very wide credible intervals were associated with these estimates. We did not identify a clear seasonal trend, however two peaks were identified, with up to 20% (95% CI = 14%, 26%) increases and 28% decreases (95% CI −35%, −22%) in risk of *R*_*c*_ associated with April/October and the beginning of January respectively (Figure 4B).

**Figure 4:**
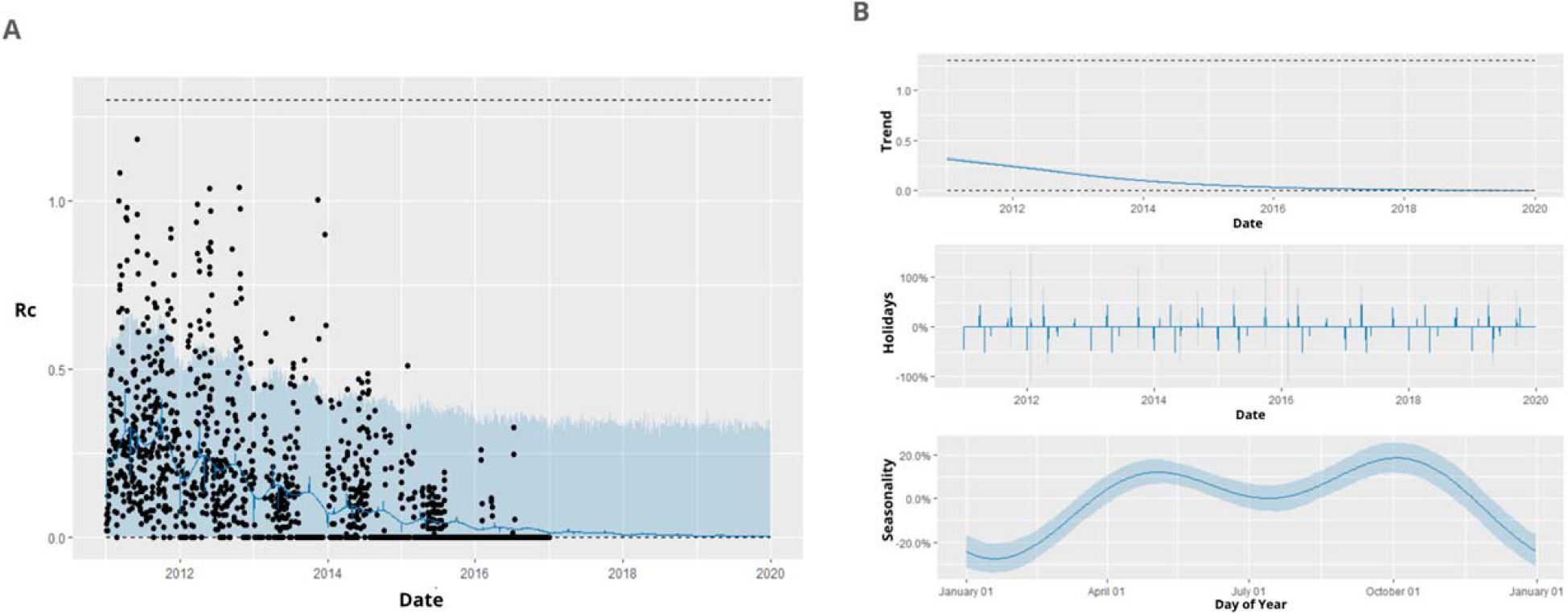
A) Black points show estimated individual *R*_*c*_ values, blue line represents prophet model predictions for mean *R*_*c*_ on that day, shaded blue area shows 95% credible interval of prediction. B) Decomposed time series model, showing the general trend, fitted holiday effect and seasonal effect. For seasonal and holiday effects the y axis shows the percentage increase or decrease in *R*_*c*_ predicted which is attributable to a seasonal or holiday effect.

## Discussion

Quantifying reproduction numbers and their spatio-temporal variation can provide useful information to inform strategies to achieve and maintain elimination in contexts where traditional measures of transmission intensity are not appropriate. We used individual level surveillance data to infer reproduction numbers by estimating the likelihood of cases being linked by transmission and applied this to a dataset of all confirmed and probable cases of *P. vivax* and *P. falciparum* occurring in Yunnan province between 2011 and 2016, which is a focus of concern for re-emergence. Our results suggest that transmission in this province decreased rapidly between 2011 and 2016 as shown by a declining risk of *R*_*c*_ exceeding zero across the province. This decline is relatively robust to assumptions about the serial interval distribution. Extrapolating this trend using time-series methods, we expect this trend to continue, predicting a mean *R*_*c*_ of 0.005 for 2019.

Given the consistently very low *R*_*c*_ values estimated by 2014 onwards, and the future projections based on observed reproduction numbers over time, our results suggest that re-emergence or outbreaks of sustained transmissions are unlikely, provided interventions are continued. However, as all data analysed was collected whilst the NMEP was in place, we cannot draw conclusions about the impact of scaling back interventions or consider other counterfactuals. There is also some uncertainty in our estimates of current and future *R*_*c*_, although the 95% credible intervals of these estimates remain below 1. It is important to note that even with low *R*_*c*_ values it is still possible for locally-acquired cases to occur following importation, however the probability of sustained chains of transmission decreases as *R*_*c*_ decreases. There also is more uncertainty in our estimates of risk in areas that have not observed many cases. It is difficult to determine whether an absence of cases is due to a lack of detection, a lack of importation events occurring or a low underlying receptivity to transmission. However, it is worth noting that the greatest uncertainty in our spatiotemporal risk estimates of *R*_*c*_ > 0 tends to be in areas of high elevation (elevation > 3000m), where there is unlikely to be transmission. Given the large numbers of imported cases, it is important to highlight these uncertainties and ensure control measures are maintained. Nonetheless, our findings are promising for China to meet their 2020 elimination goal. Our results highlight the success the country has had in malaria control and highlights the difficulty of elimination certification in contexts where both distant and local cross border importation is common.

Whilst there is a clear peak in incidence of cases occurring in May (Supplementary Figure 4), the seasonality of *R*_*c*_ estimates were less clear, although there seemed to be two peaks in seasonal increases in *R*_*c*_, one occurring in March/April, and one in October. This pattern could be an artefact of human movement, with both periods associated with seasonal movement and holiday periods – the *Chunyun* period occurs in China for Chinese New Year and the holiday week of the National Day in October and is associated with intranational travel to visit family. During this time, there is often movement from cities to rural areas, and so in these contexts there may be more opportunities for infection to occur as more people are exposed to bites from suitable vectors. This is supplemented by our finding that these specific holidays are associated with small to moderate increases in *R*_*c*_, however it is worth noting the very wide credible intervals and the great deal of uncertainty associated with these estimates, and therefore caution is required when interpreting this finding.

There are several limitations to our study. Firstly, there is a limitation in the classification of local and imported cases used in this study. For instance, the definition of importation used in case classification is defined by travel to any malaria-endemic areas outside China in the month prior to illness onset. This definition might include people who travelled abroad within the week prior to illness onset, but biologically their infection could not have been obtained during that time given the incubation period. However, in the absence of alternative information, travel history may provide a better indication of the likely importation status of a case than attempting to infer importation without this information, however there could be scope in future work to allow for incorrect travel history. As certification of elimination is now tolerant of introduced (first generation imported-to-local transmission) but not indigenous (second generation local-to-local transmission) cases, being able to differentiate between the two, and understanding how much transmission is indigenous versus imported or introduced is an important area of focus for future work.

It is important to consider unobserved cases and their potential contribution to transmission dynamics. We do account for unobserved cases via epsilon edges; however, this method is still more suited to scenarios where the majority of cases are observed. In contexts with a high level of asymptomatic infection contributing to transmission or with poor case detection and/or reporting, these approaches would not be suitable.

A second limitation is the type of data available for inference. Although not available for this study, there are several data sources that increasingly are being collected and could enhance similar analyses in the future in eliminating and pre-eliminating contexts. Firstly, methods to make use of contact tracing data have been developed for emerging outbreaks^23^ but have not to our knowledge been applied to endemic disease in the elimination. Although contact tracing for indirectly transmitted diseases is more difficult, identifying if the likely source of infection is a breeding site near the home or a place of work is carried out through active case detection schemes, but often the resulting data are not made available alongside line list data. This information could be used to weight certain connections. Genetic data are also increasingly available, and provide useful information about movement of parasites^24,25^, the likelihood of two cases being linked by transmission, and can provide useful information to help distinguish imported from local cases and chains of transmission resulting from importation from on-going local transmission^21^. Such data were not available in this context; however, a similar methodological framework or approach could incorporate information such as genetic distance. Historical data on incidence at fine scale (e.g. village level) could also be used to inform likelihood of asymptomatic infection.

We introduced a new framework for analysing individual level surveillance data and found that in Yunnan province, *R*_*c*_ has seen a notable downward trend since 2011 and is expected to remain low with maintained interventions into 2020. This decline coincides with 1-3-7 strategy in improved adherence to guidelines. We predict a mean *R*_*c*_ of 0.005 for 2019, however even with such low *R*_*c*_ values estimated, there may still be a need to continue to invest in detecting and rapidly responding to imported cases due to the large amount of human movement in order to achieve three consecutive years of zero cases and prevent resurgence. Nevertheless, China’s elimination strategy and investment in surveillance provides a useful roadmap for other countries planning for malaria elimination by illustrating how coordinated and timely surveillance and response can be implemented, as well as sustained investment in surveillance, and region-focused international collaboration.

## Methods

### Data

Anonymised case data for all confirmed (N=4078) and probable (N=285) malaria cases reported between 2011 and 2016 in Yunnan Province (N =4390) were obtained from the Chinese Centre for Disease Control (CCDC). For each case, our data included date of symptom onset, GPS coordinates of symptom onset address, health facility address, travel history, and in some cases, the GPS coordinates of presumed location of infection.

Of these cases, the majority were *P. vivax* (N = 3469, of which 2858 were classified as imported). Of all recorded *P. falciparum* cases (N=791), 91% (N=720) were imported. Small numbers of *P. malariae* (N=8) and *P. ovale* (N=1) were excluded from our analysis. Cases defined as “untyped” (N=67) were also excluded. A small number (N=27) of cases classified as mixed infection were included in the separate analyses of each species. A full breakdown of the cases and species composition across China and in Yunnan province between 2011 and 2016 is included in Supplementary Tables 1-4.

### Surveillance system in China

China has a sophisticated malaria surveillance system, described in detail elsewhere^7,15,16,26,27^ and in Supplementary Note 1. Briefly, surveillance is carried out in both a passive and reactive manner, organised and administered at the national, provincial and county level. The centralised China Information System for Disease Control and Prevention (CISDCP) receives daily updates on case reports from health facilities. The “1-3-7” strategy introduced in 2010 aiming for case reporting within one day of detection, initial investigation within three days and focused investigation and action taken in under seven days has increasingly been achieved – the proportion of cases investigated within three days increased from roughly 55% in 2011 to almost 100% by 2013. However, the programme took longer to achieve the seven day focal point investigation goals, with just over 50% of foci investigated and treated within seven days by the end of 2013^7^. Nevertheless, by 2015, adherence to the 1-3-7 strategy improved and this figure increased to an estimated 96%^27^.

### Defining the serial interval distribution

The serial interval is defined as the time between a given case showing symptoms and the subsequent cases they infect showing symptoms^28^. For a given individual *j* at time *t*_*j*_, showing symptoms before individual *i* at time *t*_*i*_, the serial interval distribution specifies the normalised likelihood or probability density of case *i* infecting case *j* based on the time between symptom onsets, *t*_*i*_ − *t*_*j*_. The serial interval summarises several distributions including the distribution of a) the times between symptom onset and infectiousness onset, b) the time for humans to transmit malaria parasites to mosquito vectors, c) the period of mosquito infectiousness, and d) the human incubation period.

Taking a similar approach to our previously developed work^20^, we defined a prior distribution of possible serial interval distributions for malaria. The serial interval distribution of treated, symptomatic *P. falciparum* malaria, previously characterised using empirical and model based evidence^29,30^ was adapted to inform the prior distribution for the relationship between time and likelihood of transmission between cases in China. We analysed *P. vivax* cases and *P. falciparum* cases separately. The prior distribution was defined to be flexible enough to reflect both the biology of *P. vivax* and *P. falciparum* as well as the dominant vector species in Yunnan (recent surveys in Yunnan province have found *Anopheles sinensis* to be the dominant vector species in mid-elevation areas and rice paddies and *Anopheles minimus* s.l. the dominant species in low elevation areas^13,31^) and to allow for possible variation in transmission dynamics, for example due to untyped infections or delays in seeking treatment. In addition, there is a possibility of a small number of asymptomatic or undetected and therefore untreated infections contributing to ongoing transmission, which will typically have a longer serial interval. We use a shifted Rayleigh distribution to describe the serial interval of both species, which can be varied by changing two parameters: *α* and *γ*. The parameter γ governs the overall shape of the distribution, and *γ* is the shifting parameter accounting for the incubation period between receiving an infectious bite and the onset of symptoms (Figure 1A). The *γ* shifting parameter was fixed at 15 days to account for the extrinsic incubation period within the mosquito and the minimum time between infection and suitable numbers of gametocytes in the blood to lead to symptom onset^32^. The prior for the *α* parameter determining the shape of the distribution was given a Normal distribution with mean 0.003 and standard deviation 0.1 (illustrated in figure 5), giving an expected time between symptom onset of one case and symptom onset of the case it infects of 36 days, with the parameter value in the 2.5 percentile of prior having an expected serial interval of 21 days and the equivalent parameter from the 97.5 percentile having an expected serial interval of 60 days. By comparison the expected values for treated *P. falciparum* from existing literature range between 33 and 49.1 days (95%CI = 33-69)^29,30^. Depending on how much uncertainty there is in the serial interval of malaria, the prior for α, the shaping parameter for the SI of malaria, may be varied. We explored the effects of different priors on the likelihood and posterior estimates. We used the same mean value for α (0.003) but set the α prior to standard deviation between 1 and 0.01. The results of considering different priors for α, the parameter shaping SI distribution on estimated *R*_*c*_ values over time is shown in Supplementary Figure 2.

**Figure 5:**
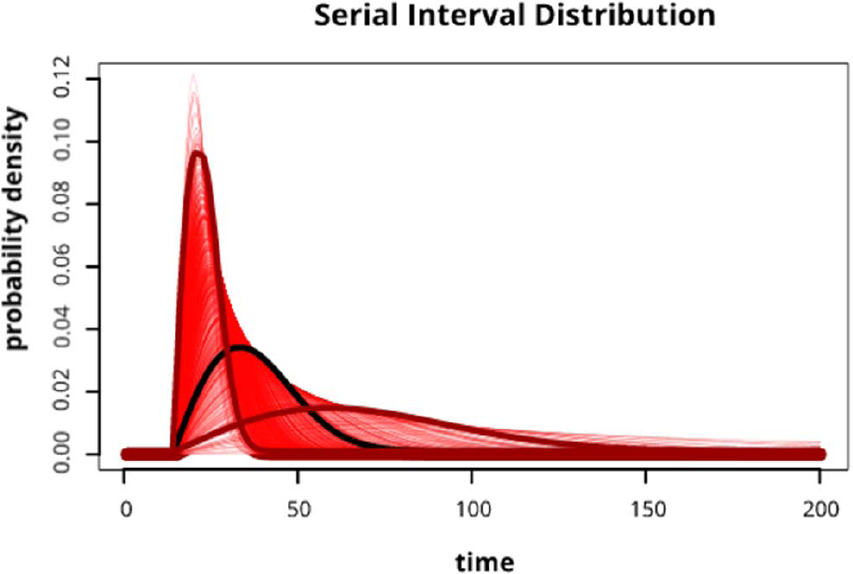
Red lines show 300 draws from the prior distribution used in the analysis for the Serial Interval distribution. The black line represents the expected function and the maroon lines represent the 2.5 and 97.5 quantile values of the prior distribution for the shaping parameter, α.

### Determining the transmission likelihood

We assume cases were classified correctly from case investigation as imported or locally-acquired based on recent travel history. Following this assumption, locally-acquired cases could have both infected others and been infected themselves. However imported cases could only infect others, as we assume their infection was acquired outside of the country. Given the evidence^7,15,27^ 2014; Hu et al., 2016; Zhou et al., 2015of strong adherence to the 1-3-7 policy for reporting and response to case detection, and no evidence of relapse within the dataset (as each patient is given a unique identifier), we assume that an individual can only be infected once by a case that has shown symptoms earlier in time.

### Transmission model specifics

To estimate the underlying pathways of transmission and likelihood of cases being linked by infection, we adapt and extend the NetRate algorithm^22^. Our adapted model introduces the ability to model serial interval functions, account for imported vs local infections and provides provision for missing or unobserved sources of infection (called epsilon edges^20,33^). We also extended the NetRate algorithm from a frequentist to a Bayesian framework to incorporate prior knowledge about the serial interval of malaria.

Consider a set of *n* infections/nodes ***I*** ∈ (*I*_1_,…, *I*_*n*_) with associated times ***t*** = {*t*_*1*_,… *t*_*n*_} ∈ ℝ^+^ and binary yes/no importation status ***π*** = {*π*_1_,.., *π*_*n*_} ∈ {1,0}. The serial interval distribution of malaria, defining the probability individual *I*_*j*_ infected individual *I*_*i*_ at times *t*_*i*_ *> t*_*j*_ is defined through a shifted Rayleigh distribution 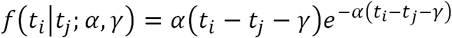 for shaping parameters *α* and *γ* (Routledge et al., 2018). For our analysis we fix *γ* = 15 days. If we assume that infections are conditionally independent given the parents of infected nodes then the likelihood of a given transmission chain can be defined as

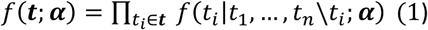

Where ***α*** is a parameter matrix. Computing the likelihood of a given transmission chain thus involves computing the conditional likelihood of the infection time of each infection (*t*_*i*_) given all other infections (*t*_1_,…, *t*_*n*_*\t*_*i*_). If we make the assumption that a node gets infected once the first parent infects it^34^ and define a survival function

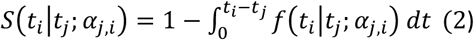

as the probability that infection *I*_*i*_ is *<u>not</u>* infected by infection *I*_*j*_ by time *t*_*i*_ then we can simplify our transmission likelihood as

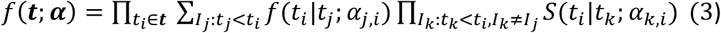

In this conditional likelihood the first term computes the probability the *I*_*j*_ infected *I*_*i*_ and the second term computes the probability that *I*_*i*_ was not infected by any *other* previous infections excluding *I*_*j*_. This likelihood therefore accounts for competing infectors and finds the infector most likely to have infected *I*_*i*_. To remove the *k* ≠ *j* condition makes the product independent of *j* and results in the likelihood

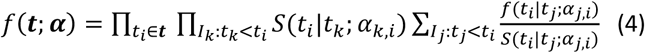

In equation 4, 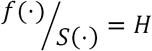 is the hazard function or rate and represents the instantaneous infection rate between individuals *I*_*i*_ and *I*_*j*_.

Assuming all cases reaching health workers or health facilities are recorded, missing cases may be generated by two processes. Symptomatic cases may be missed by not seeking care or not being found through active case detection, or cases may be asymptomatic and therefore unlikely to seek care or be detected. The latter may have densities of parasites in their blood which are too low to be detectable by microscopy if active case detection occurs. These processes apply to both imported cases or locally acquired cases. We assume the pool of asymptomatic cases in the country is low and has a small contribution to ongoing transmission. To account for unobserved infectors within our framework we include a time-independent edge that can infect any individual. The survival and hazard functions for this edge are defined as 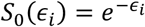 and *H*_0_ = *ϵ*_*i*_. As we will see below, as a consequence of our optimisation problem these edges are encouraged to be sparse and only invoked if no other infectors can continue the transmission chain.

In addition to unobserved edges, we assume that observed imported infectors can infect other cases but cannot be infected themselves. The final likelihood incorporating these two modifications becomes

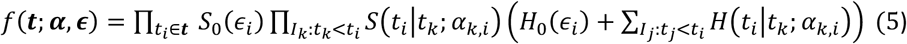

In order to find the optimal parameters for ***α***,***ϵ*** we minimize the following log likelihood subject to positive constraints on the parameters:

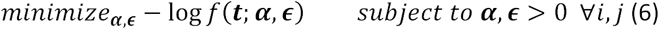

This optimisation problem is convex and guarantees a consistent maximum likelihood estimate^22^

To prevent biologically implausible serial interval distributions we impose a truncated normal prior probability distribution on ***α*** ∼Normal(0.003,0.1) [0,0.01]. When optimising our likelihood we include this prior probability and therefore evaluate the Bayesian Maximum-a-Posteriori estimate.

### Estimating *R*_*c*_

We can establish individual reproduction numbers for each case by creating a matrix where each column represents a potential infector and the rows represent a potential infectee, describing which infector edges are connected to infectees and the normalised likelihood of the cases being connected by a transmission event. Intuitively then, by taking the row sums we get the (fractional) number of secondary infections and therefore a point estimate of the time varying reproduction number *R*_*c*_(*t*_*j*_) This reflects for an individual, how many people they subsequently infect. When multiple individuals have been infected at a given time and/or place, we can take the mean individual *R*_*c*_ and uncertainty in this value as an indicator of reproduction numbers for a given time and/or location.

In contrast to traditional methods based on Wallinga and Teunis^35^ using our method in this way encapsulates not only the pairwise likelihood of transmission between two cases, but conditions this likelihood on the impact of competing edges in the inferred network (the survival of an edge). Our estimates of *R*_*c*_ therefore consider the overall transmission tree in optimisation and allow for missing cases within the tree.

### Estimating timelines towards elimination and risks of resurgence

It is important for national malaria control programmes to have information about likely timelines to elimination, chances of resurgence and uncertainty in these estimates. Using the distribution of ℛ_*c*_ values and their seasonal and general trends, we analysed time series using the *Prophet* tool and R package^36^ to explore general and seasonal trends as well as the impact of holidays on results.

This approach applies an additive regression model

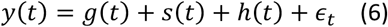

which is composed of trend, seasonal and holiday functions, where *y*(*t*) is the observations at time *t, g*(*t*) is the general trend, modelled by a logistic growth model, *s*(*t*) is the seasonal effect, modelled by Fourier series, *h*(*t*) is the effect of specific holiday dates and *ϵ*_*t*_ is the error term.

We explored the overall trend as well as seasonal trends, exploring the predicted *R*_*c*_ between 2011 and the beginning of 2020. We also explored the impact of the national holiday periods, some of which involve large scale movement, such as the *Chunyun* period around the spring festival. We cross-validated predictions and calculated root mean squared error (RMSE) and mean absolute error (MAE) (Supplementary Figure 7).

### Mapping *R*_*c*_

Transmission risk map estimates were constructed by separating individual reproduction numbers into those above and below *R*_*c*_ = 1 The latitude and longitude of the reproduction numbers were included in a binomial Gaussian random field model implemented in rINLA ^37^, in which demographic and environmental covariates (Supplementary Table 5) were used to estimate the likelihood of a case having *R*_*c*_ > 0 in the area each year from 2011 to 2016. This is a measure of malaria “receptivity” or underlying transmission potential rather than overall malaria risk, as importation likelihood is not quantified in this analysis. Area under the curve (AUC) scores from leave-one-out cross validation were used to assess model fit (Supplementary Figure 8).

## Data availability

The datasets analysed during the current study are not publicly available as they belong to the Chinese Centre for Disease Control rather than the authors but are available from Zhongjie Li on reasonable request and with permission of China CDC.

## Figure legends

Figure 3: Boxplots showing *R*_*c*_ estimates for *P. vivax* (A and B) and *P. falciparum* (C and D), aggregated by year (A and C) and month (B and D) of symptom onset. Points represent individual *R*_*c*_ estimates. Boxplots show median, upper and lower quartiles for *R*_*c*_ each.

Figure 4: Map of *R*_*c*_ estimates by year for A) *P. vivax* and B) *P. falciparum*. Blue points represent locally acquired cases; red points represent imported cases. The diameter of the point represents the size of the *R*_*c*_ estimate.

Figure 3: Map of risk of *R*_*c*_ > 0 and uncertainty in this estimate for A) *P. vivax* and B) *P. falciparum* malaria across Yunnan province in each year 2011-2016.

Figure 4: A) Black points show estimated individual *R*_*c*_ values, blue line represents prophet model predictions for mean *R*_*c*_ on that day, shaded blue area shows 95% credible interval of prediction. B) Decomposed time series model, showing the general trend, fitted holiday effect and seasonal effect. For seasonal and holiday effects the y axis shows the percentage increase or decrease in *R*_*c*_ predicted which is attributable to a seasonal or holiday effect.

Figure 5: Red lines show 300 draws from the prior distribution used in the analysis for the Serial Interval distribution. The black line represents the expected function and the maroon lines represent the 2.5 and 97.5 quantile values of the prior distribution for the shaping parameter, α.

## Acknowledgements

We are grateful to the staff members at county, prefecture, and provincial level of the Chinese Centers for Disease Control and Prevention for providing assistance with field investigation, administration and data collection in China. IR is supported by a Wellcome Trust Four Year PhD Programme grant. SL is supported by the grants from the National Natural Science Fund of China (No. 81773498), the Ministry of Science and Technology of China (2016ZX10004222-009), and the Program of Shanghai Academic/Technology Research Leader (18XD1400300). AJT is supported by funding from the Bill & Melinda Gates Foundation (OPP1106427, 1032350, OPP1134076, OPP1094793), the Clinton Health Access Initiative, the UK Department for International Development (DFID) and the Wellcome Trust (106866/Z/15/Z, 204613/Z/16/Z). KEB is funded by the Bill and Melinda Gates Foundation (OPP1197730). JLP would like to thank Bill and Melinda Gates for their active support of the Institute of Disease Modeling and their sponsorship through the Global Good Fund. The sponsors of the study had no role in the study design, data collection, data analysis, data interpretation, writing of the report, or the decision to publish. The views expressed here are those of the authors and do not necessarily represent the policy of institutions with which the authors are affiliated.

## Author contributions

**IR** Conceived idea and designed analysis and methodology, carried out analysis, wrote the original draft manuscript **SL and ZL** contributed to analysis design, collated data, interpreted the findings and commented on draft **SB** designed analysis and methodology, commented on draft, provided supervision **KEB** designed map visualisation, commented on draft **ACG** commented on draft, contributed to analysis design, provided supervision, **KBG and JLP** contributed to methodology design and commented on draft, **MGR** designed methodology **SM and AJT** commented on draft

## Materials & Correspondence

For questions regarding data access and collection, please contact Zhongjie Li. For all other enquiries please contact the corresponding author.

## Ethics approval and consent to participate

It was determined by the National Health and Family Planning Commission, China, that the collection of malaria case reports was part of continuing public health surveillance of a notifiable infectious disease. The ethical clearance of collecting and using second-hand malaria data from the surveillance was also granted by the institutional review board of the University of Southampton, UK (No. 18152). All data were supplied and analysed in an anonymous format, without access to personal identifying information.

