## Supplementary Information for "Tracking progress towards malaria elimination in China: estimates of reproduction numbers and their spatiotemporal variation"

### **Contents**

##### Supplementary Figure 1: Histogram of $R_{c}$ estimates

##### Supplementary Figure 2: $R_{c}$ over time for *P. falciparum* in Yunnan province.

##### Supplementary Figure 3: $R_{c}$ over time for *P. vivax* in Yunnan province.

##### Supplementary Figure 4: $R_{c}$ estimates by month compared to number of cases observed by month

##### Supplementary Figure 5: Histogram of epsilon edge distribution

##### Supplementary Figure 6: Map of standard deviation in estimates of P($R_{c}$ >0) for A) *Plasmodium falciparum* and B) *Plasmodium vivax*

##### Supplementary Figure 7: Cross validation of timeseries analysis

##### Supplementary Figure 8: ROC curve plot for map of P($R_{c}$ >0)

##### Supplementary Note 1: Surveillance system in China

##### Supplementary Table 1: Cases by diagnosis type (probable and confirmed) and species across China

##### Supplementary Table 2: Cases by imported/local status and species across China

##### Supplementary Table 3: Cases by diagnosis type (probable and confirmed) and species across Yunnan Province

##### Supplementary Table 4: Cases by imported/local status and species across Yunnan province

##### Supplementary Table 5: Environmental and demographic covariates used

##### Supplementary Figure 1: Histogram of $R_{c}$ estimates for A) *P. vivax* in Yunnan and B) *P. falciparum* in Yunnan. Dotted lines show median, solid lines show mean.

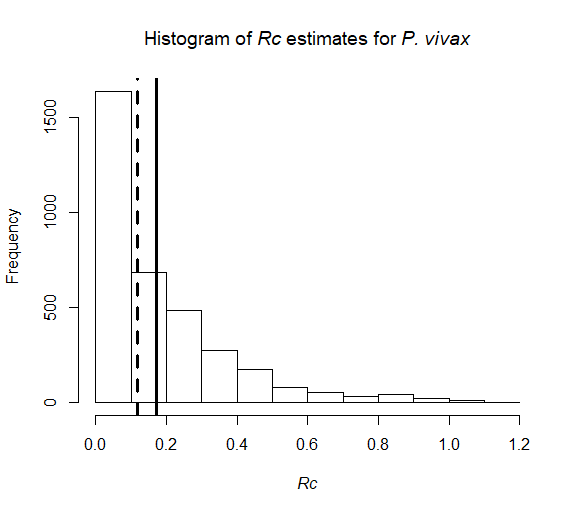

**A**

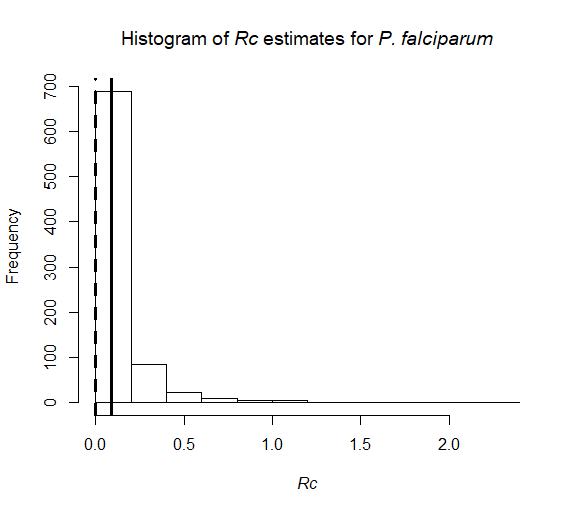

**B**

##### Supplementary Figure 2: Maximum a posteriori estimates of$R_{c}$over time for *P. falciparum* in Yunnan province. Colours represent different standard deviations of prior for α, the shaping parameter for the serial interval distribution.

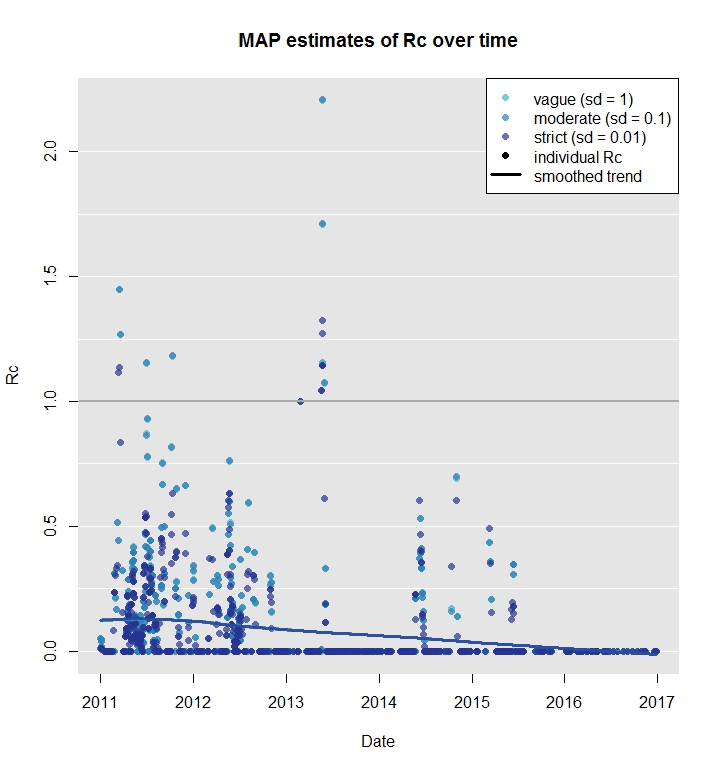

##### Supplementary Figure 3: $R_{c}$ over time for *P. vivax* in Yunnan province. Colours represent different standard deviations of prior for α, the shaping parameter for the serial interval distribution.

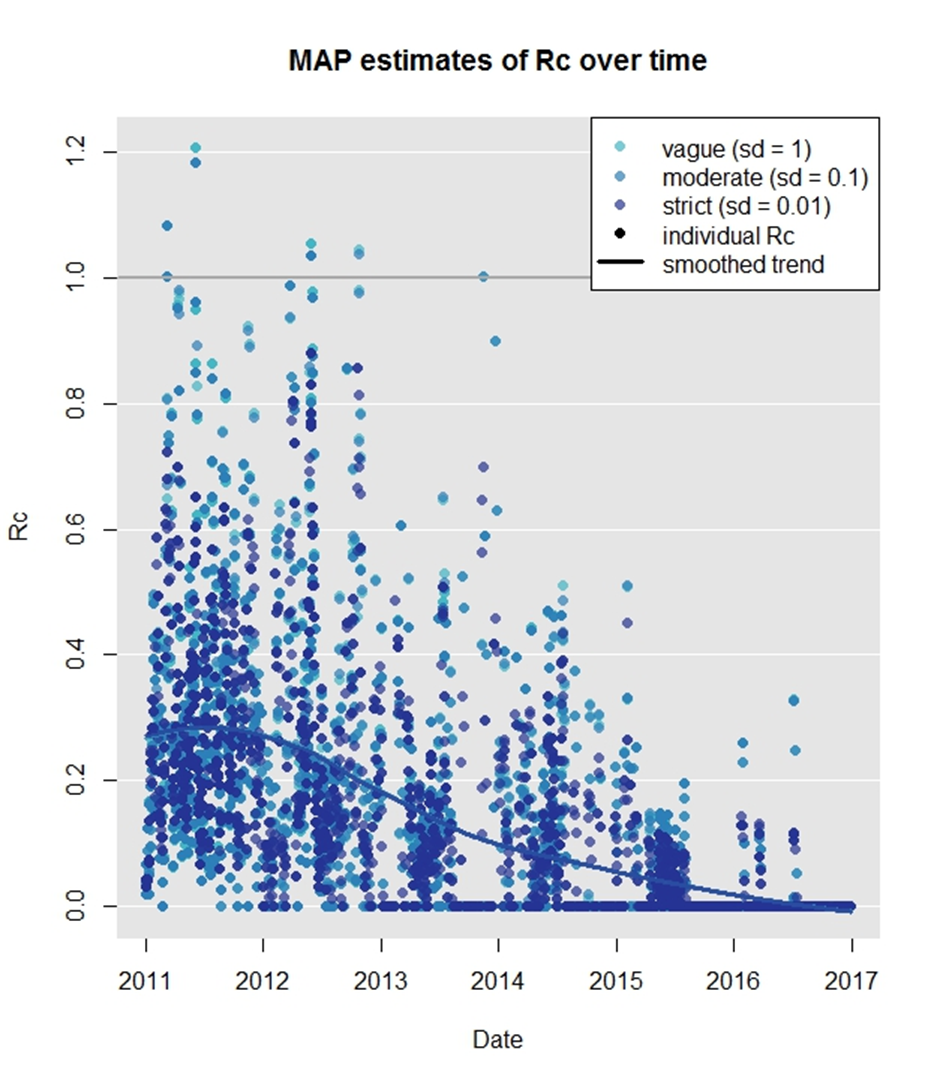

##### Supplementary Figure 4: $R_{c}$ by month compared to cases. A) overall B) stratified by year-month C) stratified by month-year

**A**

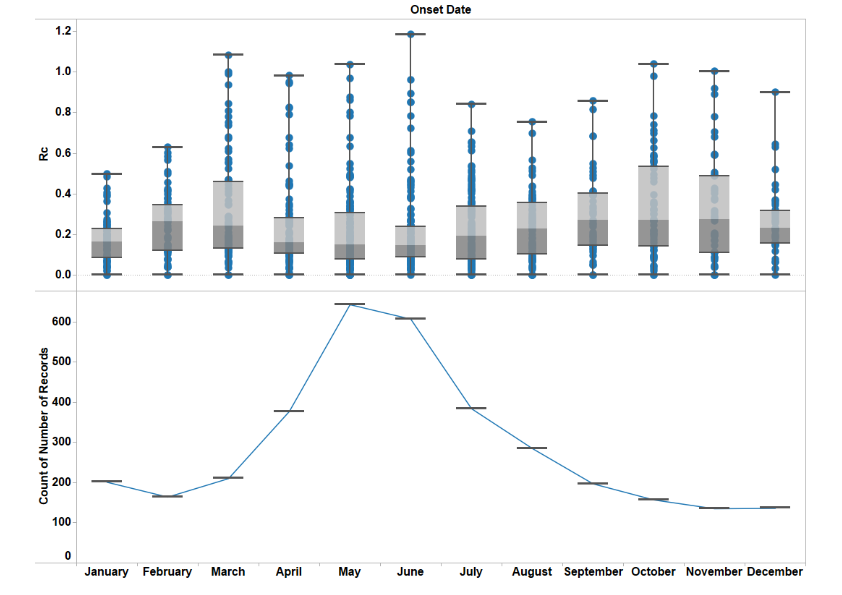

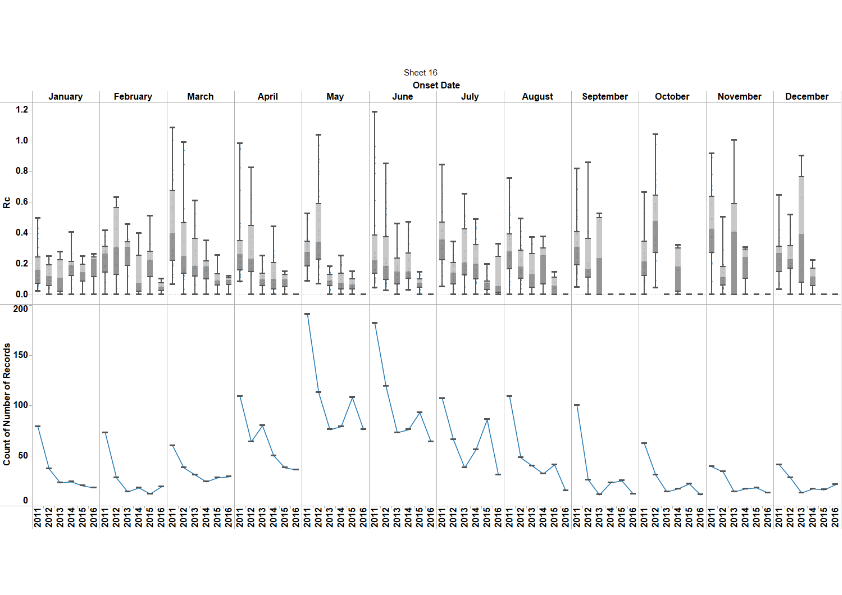

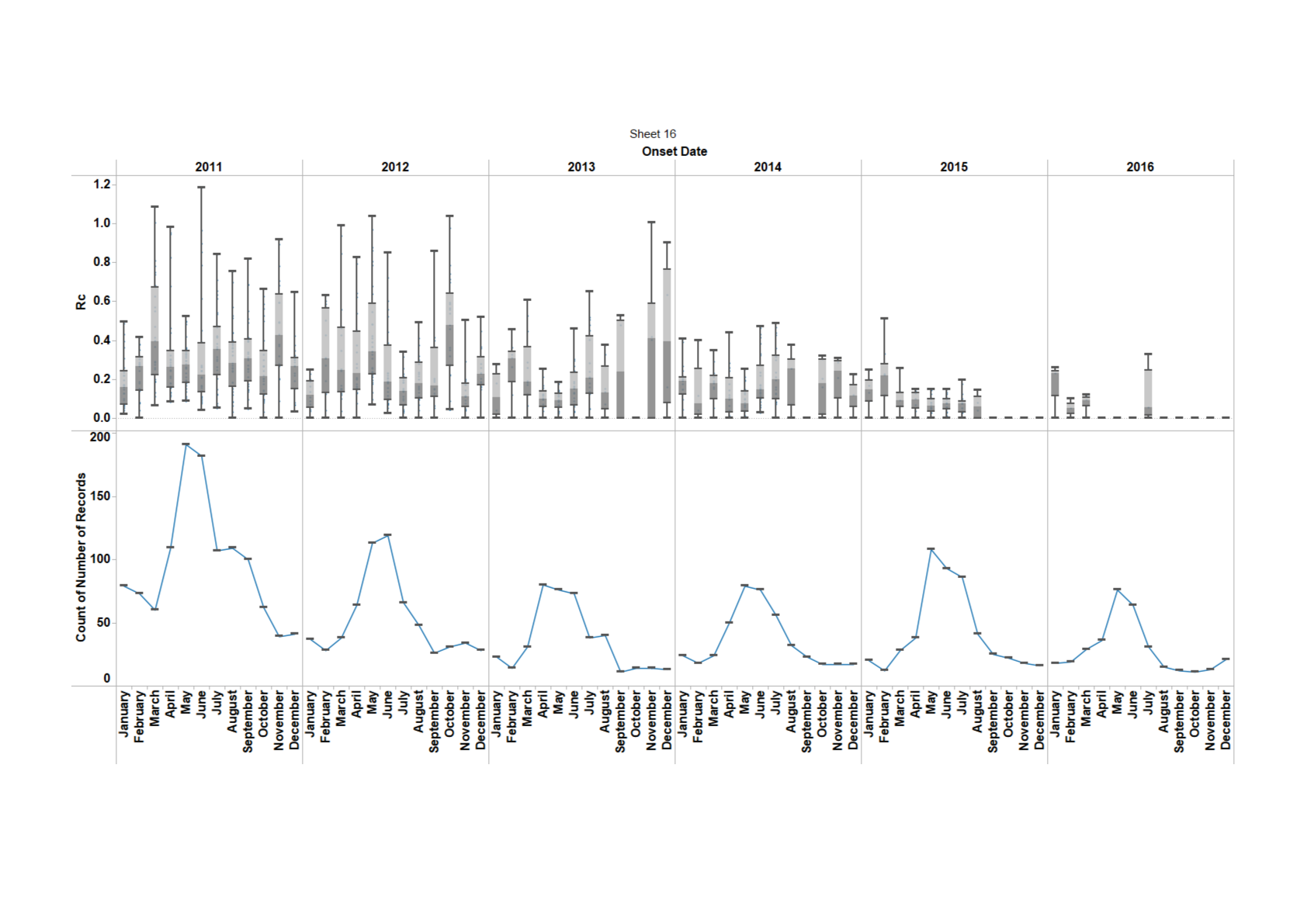

**B**

**C**

##### Supplementary Figure 5: Histogram of epsilon edge distribution for A) *P. vivax*, B) *P. falciparum*

**A**

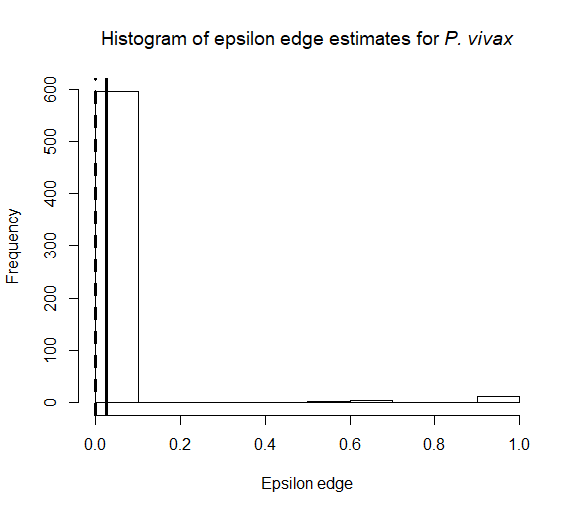

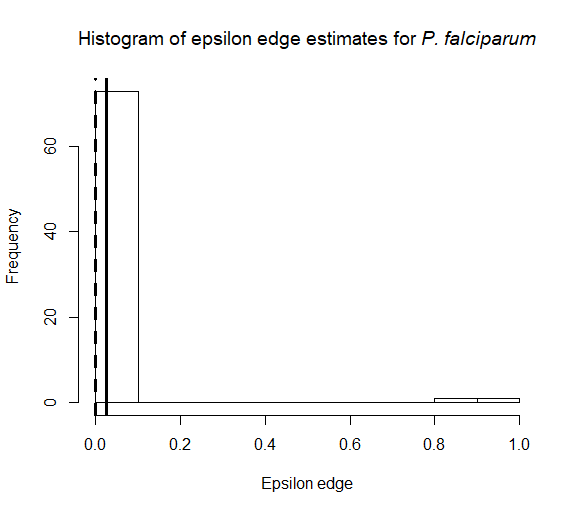

**B**

#### Supplementary Figure 6: Map of standard deviation in estimates of P($R_{c}$ >0) for A) *Plasmodium falciparum* and B) *Plasmodium vivax*

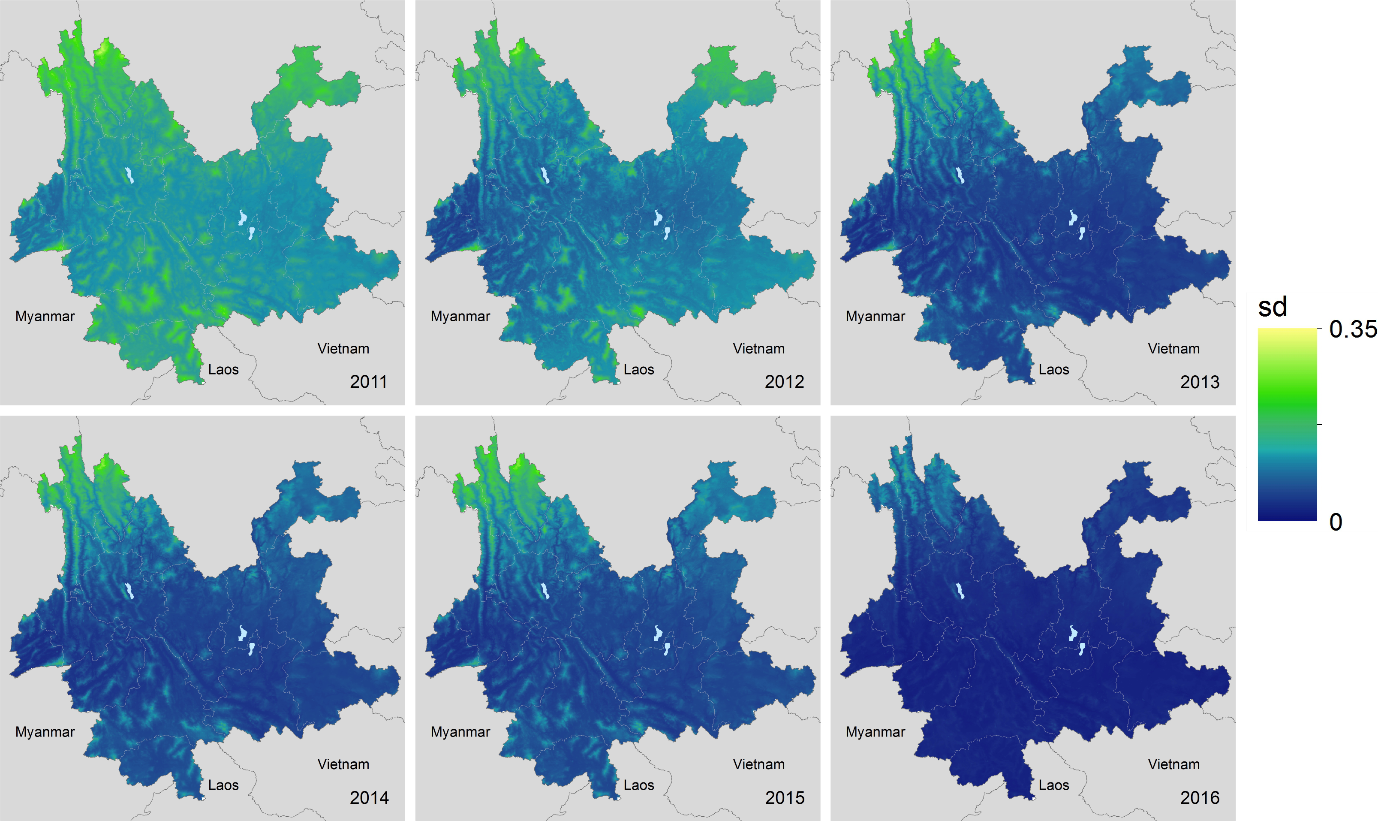

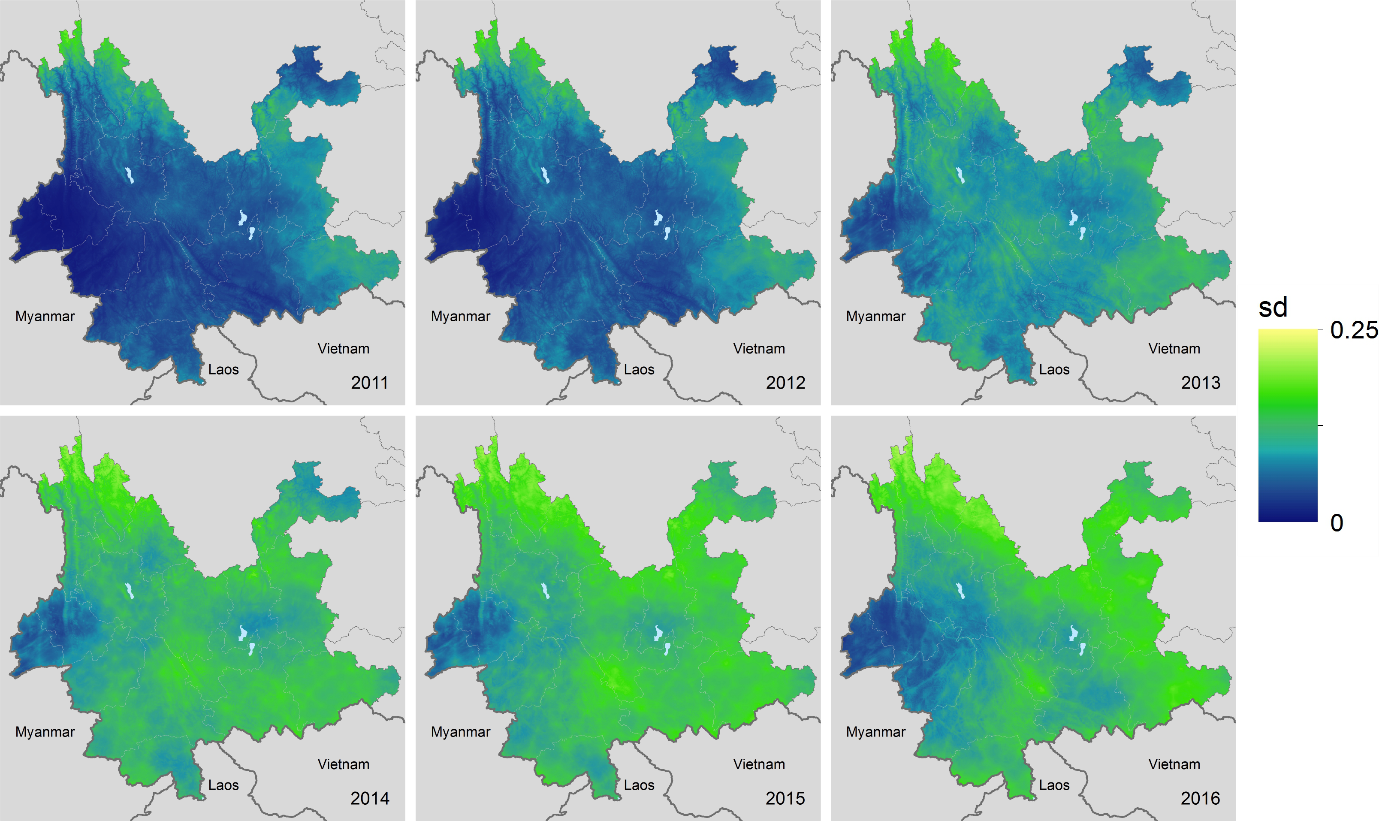

##### Supplementary Figure 7: Cross validation of timeseries analysis

The training set used was the first 730 days of data and the horizon used was 365 days.

**MAE**

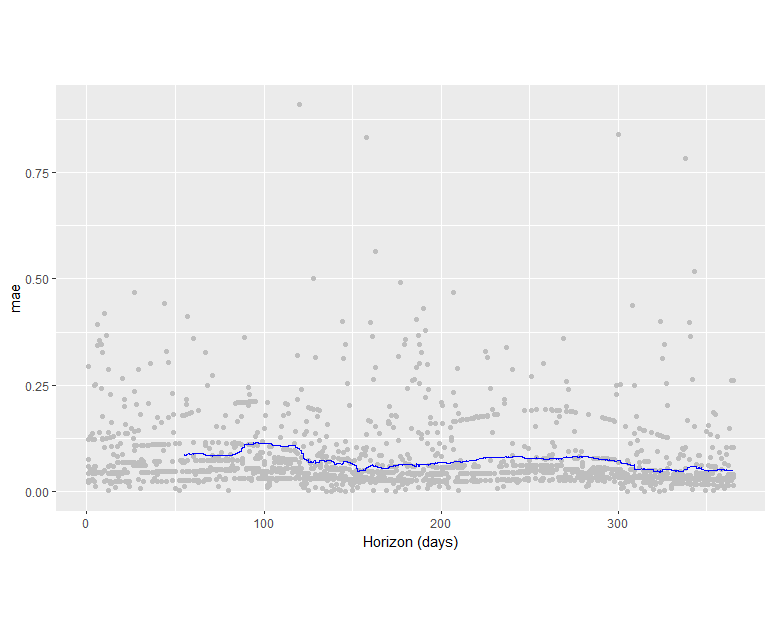

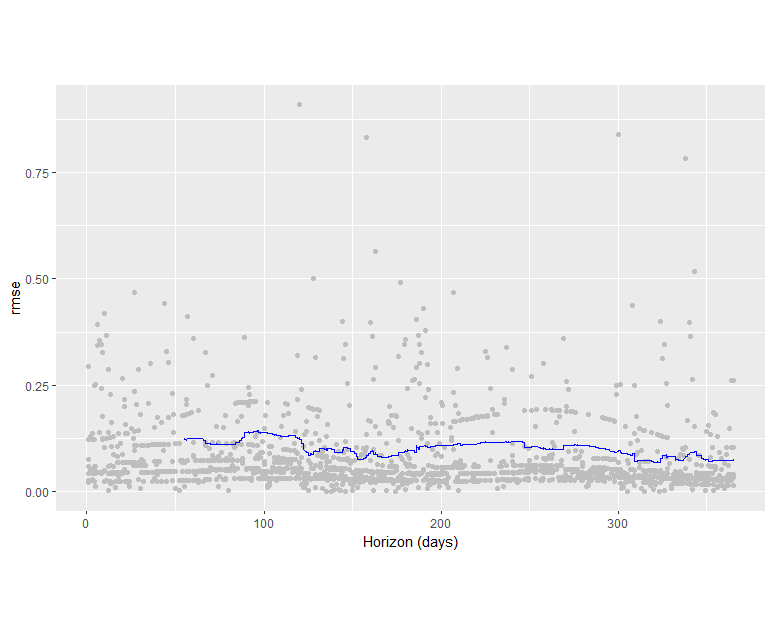
**RMSE**

##### Supplementary Figure 8: ROC curve plot for map of P($R_{c}$ risk >0) for A) *P. vivax* B) *P. falciparum*

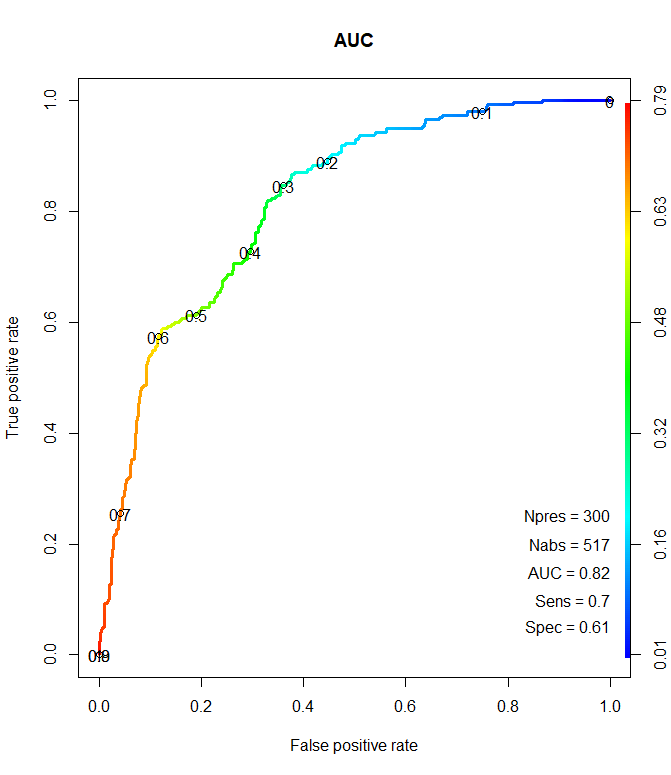

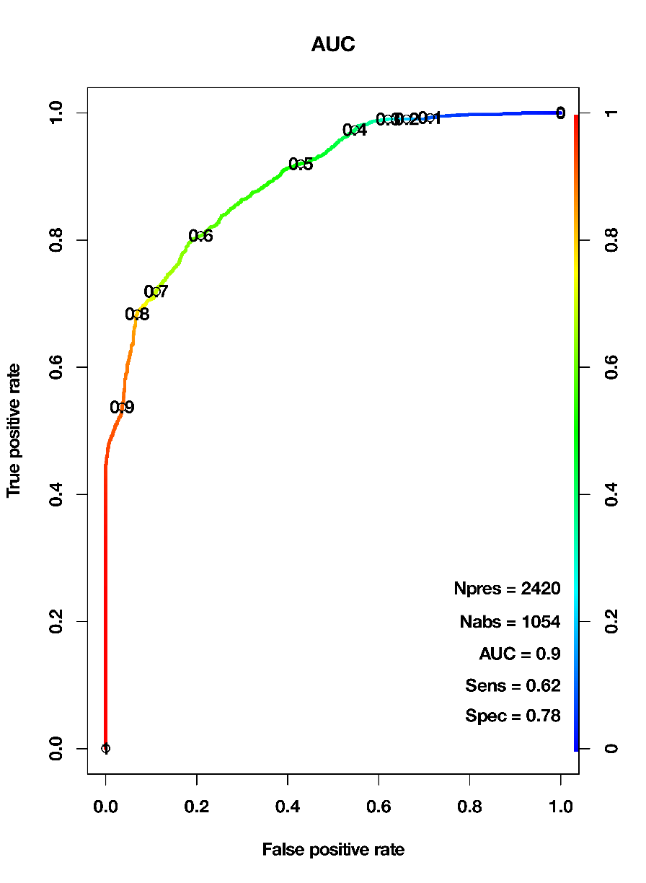

**B**

**A**

##### Supplementary Note 1: Surveillance system in China

The PRC has a sophisticated malaria surveillance system, described in detail elsewhere ^12–16^. Briefly, surveillance is carried out in both a passive and reactive manner, organised and administered at the national, provincial and county level. The centralised China Information System for Disease Control and Prevention (CISDCP) receives daily updates on case reports from health facilities

Passive detection occurs according to a protocol at the local level, such that cases are tested by microscopy or Rapid Diagnostic Test (RDT) and reported to the central information system within 24 hours. Case investigation is then pursued, where cases are confirmed via double readings of microscopy slides and in some cases polymerase chain reaction (PCR) confirmation at provincial laboratories. At this point it is also determined whether the case is locally acquired or imported by taking patient travel history – if a patient has travelled to a malaria endemic country within a month of symptom onset the case is then classified as imported ^14^. Case investigation should be completed within three days.

Foci investigation occurs once a case is detected to determine whether the foci is inactive, active or a pseudo-focus based upon the absence or presence of suitable vectors (inactive), and presence or absence of malaria in the resident area of the case if imported (pseudo-focus). Reactive Case Detection (RACD) of case contacts and populations with demographic similarities (for example individuals working in the same industry and vicinity as the case) is carried out. In active foci more intensive RACD screening of a larger pool of neighbours and contacts is carried out using Rapid Diagnostic Tests (RDTs) for immediate detection, followed by PCR testing of blood spots to detect low-density infections. IRS (Indoor Residual Spraying) is also carried out^12,14,16^.

The Ministry of Health (MoH) in China has also been measuring the timeliness of the recommended protocol and follow-on ability to meet these targets. It was found that the one-day target for case reporting was almost always met because this is required by law**.** In the years following the introduction of the 1-3-7 policy, the proportion of cases investigated within three days increased from roughly 55% in 2011 to almost 100% by 2013. However the programme took longer to achieve the seven day focal point investigation goals, with just over 50% of foci investigated and treated within seven days by the end of 2013 ^14^. Nevertheless, by 2015, adherence to the 1-3-7 strategy improved and this figure increased to an estimated 96% ^16^**.** Whilst some cases could still be missed, the thoroughness of the approach means numbers of missing cases are likely to be small.

##### Supplementary Table 1: Cases by diagnosis type (probable and confirmed) and species across China

|  | **Mixed infection** | ***P. falciparum*** | ***P. malariae*** | ***P. ovale*** | ***P. vivax*** | **Untyped** |
| --- | --- | --- | --- | --- | --- | --- |
| **Confirmed** | 260 | 11830 | 252 | 822 | 6631 | 87 |
| **Probable** | 0 | 176 | 0 | 0 | 693 | 311 |

##### Supplementary Table 2: Cases by imported/local status and species across China

|  | **Mixed infection** | ***P. falciparum*** | ***P. malariae*** | ***P. ovale*** | ***P. vivax*** | **Untyped** |
| --- | --- | --- | --- | --- | --- | --- |
| **Local** | 5 | 92 | 4 | 1 | 1711 | 95 |
| **Imported** | 255 | 11914 | 248 | 821 | 5613 | 303 |

##### Supplementary Table 3: Cases by diagnosis type (probable and confirmed) and species across Yunnan Province

|  | **Mixed infection** | ***P. falciparum*** | ***P. malariae*** | ***P. ovale*** | ***P. vivax*** | **Untyped** |
| --- | --- | --- | --- | --- | --- | --- |
| **Confirmed** | 27 | 770 | 8 | 1 | 3269 | 3 |
| **Probable** | 0 | 21 | 0 | 0 | 200 | 64 |

##### Supplementary Table 4: Cases by imported/local status and species across Yunnan province

|  | **Mixed infection** | ***P. falciparum*** | ***P. malariae*** | ***P. ovale*** | ***P. vivax*** | **Untyped** |
| --- | --- | --- | --- | --- | --- | --- |
| **Local** | 4 | 71 | 0 | 0 | 611 | 51 |
| **Imported** | 23 | 720 | 8 | 1 | 2658 | 16 |

##### Supplementary Table 5: Environmental and demographic covariates used

| variable class | variable(s) source type | | |
| --- | --- | --- | --- |
| temperature | land surface temperature (day, night and diurnal-flux) | MODIS product | dynamic monthly |
| precipitation | mean annual precipitation | WorldClim | synoptic |
| elevation | digital elevation model | SRTM | static |
| infrastructural development | accessibility to urban centres and night-time lights | modelled product and VIIRS | static |
| moisture metrics | aridity and potential evapotranspiration | modelled products | synoptic |
